# Replidec - Use naive Bayes classifier to identify virus lifecycle from metagenomics data

**DOI:** 10.1101/2022.07.18.500415

**Authors:** Xue Peng, Jinlong Ru, Mohammadali Khan Mirzaei, Li Deng

## Abstract

**Motivation:** Viruses are the most abundant biological entities on earth. The majority of these entities are bacterial viruses or phages which specifically infect bacteria. Phages can use different replication strategies to invade their hosts including lytic, lysogenic, chronic cycle and pseudolysogeny. While the determination of the replication strategy used by phages is important to explore the phage-bacteria relationships in different ecosystems there are not many tools that can predict this in metagenomic data. In addition, most of the tools available can only predict lytic and lysogenic cycles. To address this issue, we have developed a new software called Replidec to identify three most common phage replication cycles (virulent, temperate, chronic) in viral sequences.

**Results:** Replidec uses Naive Bayes classifier combined with alignment-based methods to improve the prediction accuracy in metagenomic data. We test Replidec on viral genomes with known replication cycle and simulated metagenomic sequences. Replidec perform relatively good both in isolated genomes (F1 score: 92.29% ± 0.81; mcc: 89.14% ± 1.22) and simulated metagenomic sequences(F1 score: 87.55% ± 2.12; mcc: 88.23% ± 2.55). Moreover, Replidec can also accurately predict the replication cycle in small viral fragments(∼3000bp). In conclusion, Replidec can achieve the best performance in simulated metagenomic data compared to most prediction softwares including BACPHLIP.

## Background

As the most biological abundant entities on earth, the total abundant of viruses was estimated as 10^31 on the planet [1]. They regulate biogeochemical cycles in most ecosystems, including the human body, through interacting with their bacterial hosts[2, 3]. The majority of these viruses are bacteriophages or phages [2, 3]. Phage can replicate through different replication cycles, including lytic, lysogenic, chronic cycle and pseudolysogeny.

Lysogenic cycle happens when a phage enters into a stable symbiosis with its bacterial host through integrating its genome into the host chromosome, or is maintained as a plasmid and replicates with the bacterial host (the latter can be referred as pseudolysogenic cycle). The host and phage in such a relationship are called a lysogen and a prophage, respectively. Prophages can enter the lytic cycle either spontaneously at a low frequency or in response to external stressors such as those that trigger DNA damage or the SOS response[4][5][6]. Chronic cycle, as seen for filamentous phages e.g. phage M13, is a replication cycle whereby virions are produced and continuously released without killing the host[7]. The characterization of phage’s replication cycles can shed some light on the unknown phage-bacteria interactions in complex ecosystems, including the human body, and contribute to understanding the role of (pro)phages in human health and disease.

The recent advances of high-throughput shotgun sequencing and the increase in their application in virome research has enabled significant advances in the field. Shotgun sequencing bypasses the need for culturing and allows for studying viruses of complex ecosystems such as the human body. As a result, the total number of uncultivated viruses sequenced each year is far more than virus isolates [8], and uncultured viruses already represent most of the viral diversity in public databases[8]. Multiple tools have been developed to analyze these novel uncultivated viruses *in silico*, which can be used for genome annotation, taxonomic classification, and host prediction [8]. Also, there is a need to develop tools for making predictions about the replication cycle for these uncultivated viruses from viral metagenome, or virome data. The prediction of the viral replication cycle would provide valuable insights on virus-host interactions in complex ecosystems and the role of virome in human health and disease [9][10][4].

We can divide the current available tools for in silico prediction of the replication cycles in viruses into two groups: alignment based methods (such as PHACT [11], BACPHLIP [12]), and alignment free based methods (PhageAI [13], PhagePred [14], Deephage [15]). Yet, only PhagePred, Deephage are designed for virome data.

Alignment-based methods. PHACTS is a machine learning classifier that predicts phage replication cycles from the protein sequences encoded in the genome. PHACTS uses a similarity algorithm and a supervised Random Forest classifier to predict whether a phage prefers a lytic or lysogenic cycle. Yet, PHACTS was trained and tested on a small set of phages, a total of 227, may affect its prediction accuracy [11]. BACPHLIP uses a Random Forest classifier to detect the presence/absence of a set of the conserved proteins associated with lysogeny in phage genomes to predict their replication cycle. BACPHLIP was trained on 634 phage genomes and has shown 98% accuracy on a set of 423 phages. However, it requires the input phage genome to be complete to predict the phage’s replication cycle confidently[12].

### Alignment-free methods

PhageAI relies on Word2Vec with Ship-gram model and linear Support Vector Machine to predict phage’s replication cycle. PhageAI has achieved 98.90% accuracy on a validation set and it is a REST web service and available as Python package with limited to 100 requests/day [13]. PhagePred uses *k*-mer frequencies as the sequence feature to evaluate the 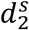 dissimilarity measures between novel viral sequences and two types of replication cycles in phages. However, the *k*-mer–based methods often generate a high level of noise with short sequences and might also lose detailed sequence information when encoding sequence to *k*-mer feature vectors[14]. DeePhage uses a convolutional neural network for detecting signatures associated with lytic/lysogenic cycle in viral sequences to predict their replication cycle. DeePhage is trained on well-annotated complete phage genomes from public databases, and its accuracy of prediction reaches as high as 89%. However, DeePhage cannot identify eukaryotic viruses within virome data before predicting their replication cycle, which can cause bias to appear in predictions [15]. All the tools mentioned can only predict lytic, and lysogenic cycles, except PhageAI that also predicts chronic.

Here, we developed a new software (Replidec) to identify the possible replication cycle (virulent, temperate, chronic) of viral sequence. The query viral sequence can be assembled contigs from metagenomic sample or genome from a isolated virus sample. Replidec use Naive Bayes classifier combine with alignment-based method to improve the prediction accuracy. We use a three step approach for predicting viruses replication cycle: 1) detecting the presence of two well-described phage proteins associated with lysogeny (integrase and excisionase) in viral contigs, 2) using a Naive Bayes classifier to calculate the conditional probability based on alignment to an in house database and 3) assigning chronic replication cycle by identify PI-like genes (https://doi.org/10.1038/s41564-019-0510-x). Replidec predicts the replication cycle of the viral contigs based on the outcomes of these three steps using a voting system (Figure 4). The code is available at Github: https://github.com/deng-lab/Replidec

## Results

### Function of Viral_Protein_DB

We create a custom viral protein database (Viral_Protein_DB) used as the training dataset in Bayes classifier. Viral_Protein_DB contain 1,323,491 CDS, and it plays a key role in prediction of phages replication cycle by Replidec. We check the function distribution of Viral_Protein_DB using 4 databases (see material) to annotate the amino acid sequence of all CDS in Viral_Protein_DB. We were able to annotate 74.59% (987,225) of most proteins using these databases while one fourth of the sequences remained unannotated (Supplement S1). The most frequent function in all annotated sequences is related to phages structural proteins such as tail protein (∼19k), minor tail protein (∼16k), portal protein (∼15k), terminase large subunit (∼15k) and more (Supplement S1). In addition, integrase (∼19k), endolysin (∼16k), transcriptional regulators (12,366) and transcriptional repressors(7,818) are also highly frequent in Viral_Protein_DB. The majority of transcriptional regulator genes (11,140 from temperate virues and 1,226 from virulent virues) and transcriptional repressor genes (7,717 from temperate virues and 101 from virulent virues) are from temperate phages. Some of these genes control the lysis-lysogeny decision, such as CI, CII and Cro in phage λ[16].

### Functional annotation of Viral_Protein_DB protein cluster

We also check the function of the high tendency protein cluster. The tendency of each protein cluster is measured by the log form of probability of lysogenic divided by the probability of Lytic. That is if the value of a protein cluster is greater than zero, the protein cluster has a tendency to lysogenic and vice versa. We only pick high tendency clusters with the following criteria: a) cluster size great than 100 and b) the tendency great than 4 or small than -5 (Supplement S2). PC_392291 (Integrase) show a high lysogenic tendency and it accordance with network analysis that lysogenic and excisonase gene are found in temperate phage sub-network(Fig 1B). PC_408056 show highest lysogenic tendency, unfortunally the function of this protein cluster is unknown. PC_67795 (endolysin, phrog_102) show a high lysogenic tendency while PC788848 (endolysin, phrog_7) show a high lytic tendency. This may demonstrate that temperate virus and virulent virus may use different lysis system to lysis host cell.

**Figure 1:**
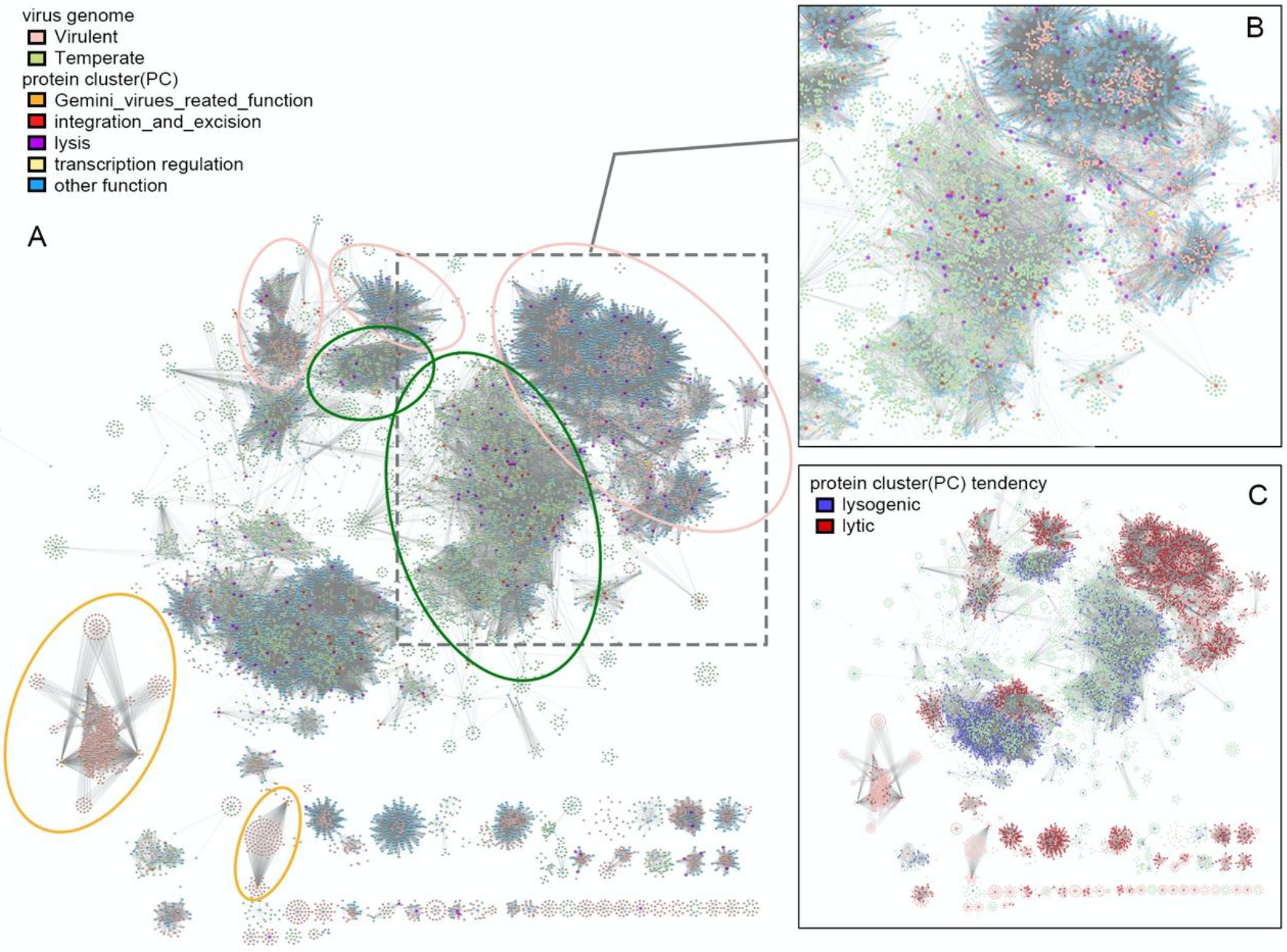
Network between protein clusters and virus genomes. (A and B) network of protein clusters colored by functions and (C) network of protein clusters colored by tendency. Light green nodes are temperate virus genomes and light pink nodes are virulent virus genomes. Protein clusters related to structure and DNA, RNA_and_nucleotide_metabolism are removed to lower the figure’s complexity.

Protein cluster with endonuclease related function show a high lytic tendency such as PC_6602 (endonuclease, phrog_755), PC_642569 (SbcD-like subunit of palindrome specific endonuclease, phrog_100), PC_555232 (endonuclease VII, phrog_423|phrog_24351), PC_273068 (DenB-like endonuclease IV, phrog_1360), PC_697996(SbcC-like subunit of palindrome specific endonuclease, phrog_77). They are key components of phage DNA packaging machines in replication cycle[17].

Unexpected, protein cluster with function related to Gemini viruses, such as PC_782405 (Gemini_AL1, Gemini_AL1_M), PC_776530(Gemini_coat), PC_585642(Gemini_coat), PC_85974(Gemini_AL3), PC_63128 (Gemini_BL1) also show high virulent tendency. From the network analysis, these protein clusters are only exists in some certain viruses genomes(Fig 1 A). These function motif is related with Geminiviruses and take part in viral replication and encapsidation. Geminiviruses have small, single-stranded DNA genomes which encode 5–7 proteins that can redirect host machineries and processes to establish a productive infection by interact with multiple plant protein kinases and hormone pathways[18].

### Gene-sharing network between protein clusters and virus genomes

To check the relationship between genomes and protein clusters, a gene-sharing network was created; the genomes of temperate and virulent viruses formed different sub-network (Fig 1). We color coded the protein cluster with function (Fig 1A) and tendency(Fig 1C), respectively. Genes with function related to integration and excision are found mostly in temperate viruses(Fig 1 B), while protein clusters associated with lysis are spread all over the network. There are also many protein clusters with unknown functions that are associated with virulent and temperate viruses sub-network. Therefore, relying only on genes with known function to predict replication cycle will not provide a clear picture of the diversity of phage replication cycle. Replidec use both known and unknown genes to increase the prediction accuracy. Our analyses show that the virulent viruses use different sets of genes showing tendency to these viruses in the network compared to temperate viruses through their replication cycle. For examples, the Geminivirues related functions show the highest lytic tendency, exist in virulent viruses, and they exist in two sub-networks(Figure1 A orange circle).

### Comparing PhageAI, Bacphlip, PHACTS and Replidec accuracy of predictions using complete genomes

We compared the performance of these tools in predicting the replication cycles of phages using complete genomes(Figure 2). PHACTS, the first tool developed for predicting the replication cycle of viruses showed lower accuracy. In contrast, PhageAI have the best performance in sensitivity (98.65% ± 0.54), accuracy (98.46% ± 0.55), F1 score (98.60% ± 0.57) and matthews correlation coefficient (mcc, 97.55% ± 0.92). But the PhageAI API have query limitations (100 query/day), so batch query for a large volume of sequences will not be possible. Replidec perform better than BACPHLIP in sensitivity, accuracy, F1 score and mcc. The average sensitivity of Replidec (92.65% ± 0.87) is higher than BACPHLIP (64.72% ± 0.62). The Replidec (F1 score: 92.29% ± 0.81; mcc: 89.14% ± 1.22) also showed a better performance (F1 score: 59.21% ± 1.14; mcc: 68.83% ± 1.63) in average F1 score and mcc.

**Figure 2.**
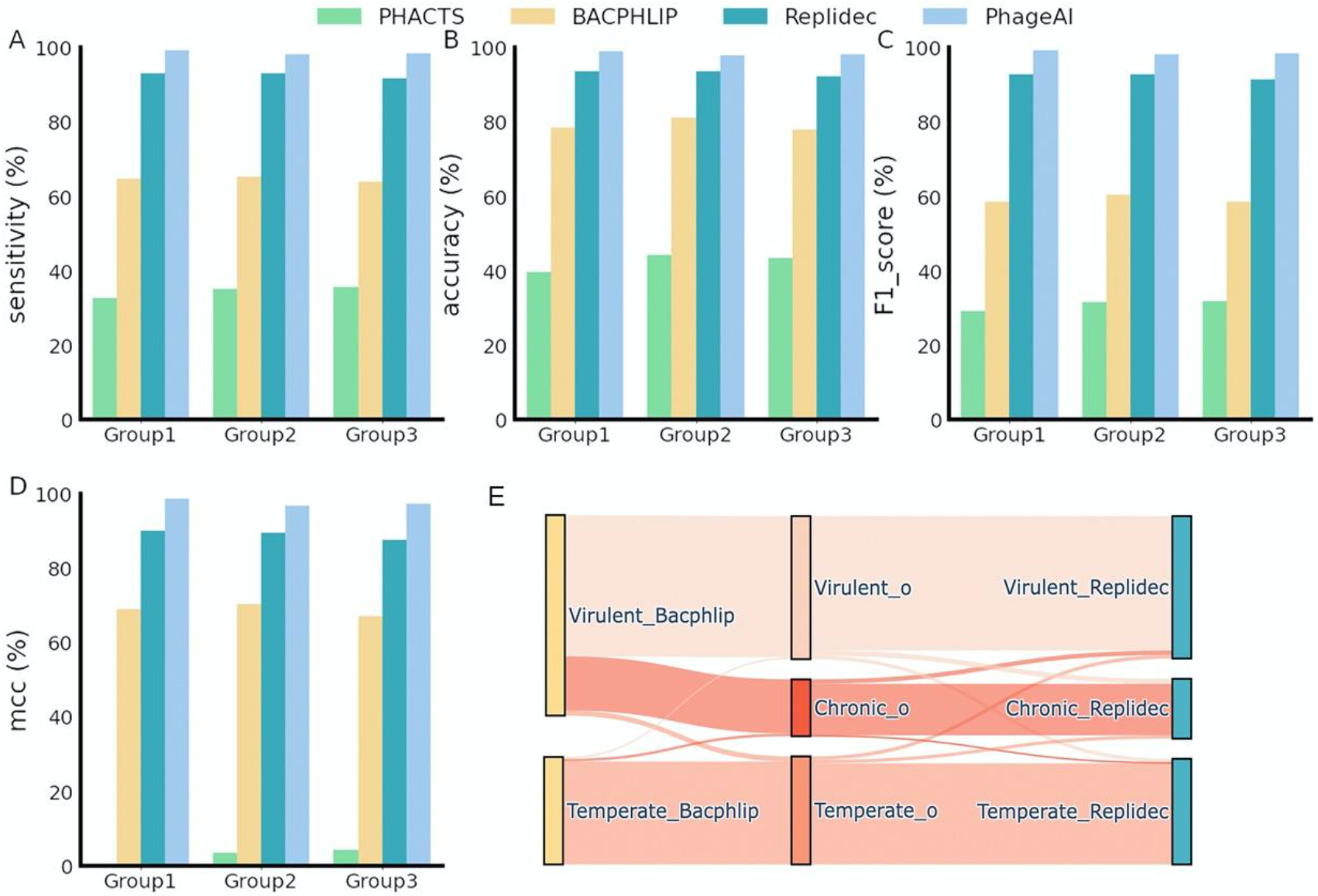
Performance of different tools in predicting life cycle of complete viral genomes. Comparison between four softwares on sensitivity(A), accuracy(B), f1_score(C), mcc(D). (E) detailed comparison between Bacphlip (left bar), Replidec (right bar) and original test data(middle bar with o as suffix).

To investigate the difference between BACPHLIP, Replidec, we use sankey plot to compare the predictions by BACPHLIP, Replidec and original data. One more limitation with BACPHLIP is that it can not predict chronic replication cycle, so most of chronic viruses are classified as virulent (Figure2 E).

### Comparing prediction accuracy of DeePhage, Bacphlip, PHACTS and Replidec using metagenomic simulated assemblies

We tested the performance of these tools on simulated assemblies (Figure 3), it should be noted that PHACTS and BACPHLIP are developed for predicting the lifestyle using complete viral genomes while DeePhage is created for prediction replication cycle for metagenomic-derived sequence. PHACTS showed the lowest accuracy among all tools tested. The average sensitivity of Replidec and DeePhage is 89.30% ± 1.06 and 59.17% ± 0.74, respectively. The average sensitivity of BACPHLIP reach to 39.89% ± 0.47 slightly higher than PHACTS(33.70% ± 0.96). Replidec show highest value among other softwares in accuracy (93.78% ± 1.41), F1 score (87.55% ± 2.12) and matthews correlation coefficient index (mcc, 88.23% ± 2.55). And DeePhage take the second place at 84.29% ± 1.06%, 57.23% ± 0.88%, 70.48% ± 2.45% in accuracy, F1 score, mcc.

**Figure 3.**
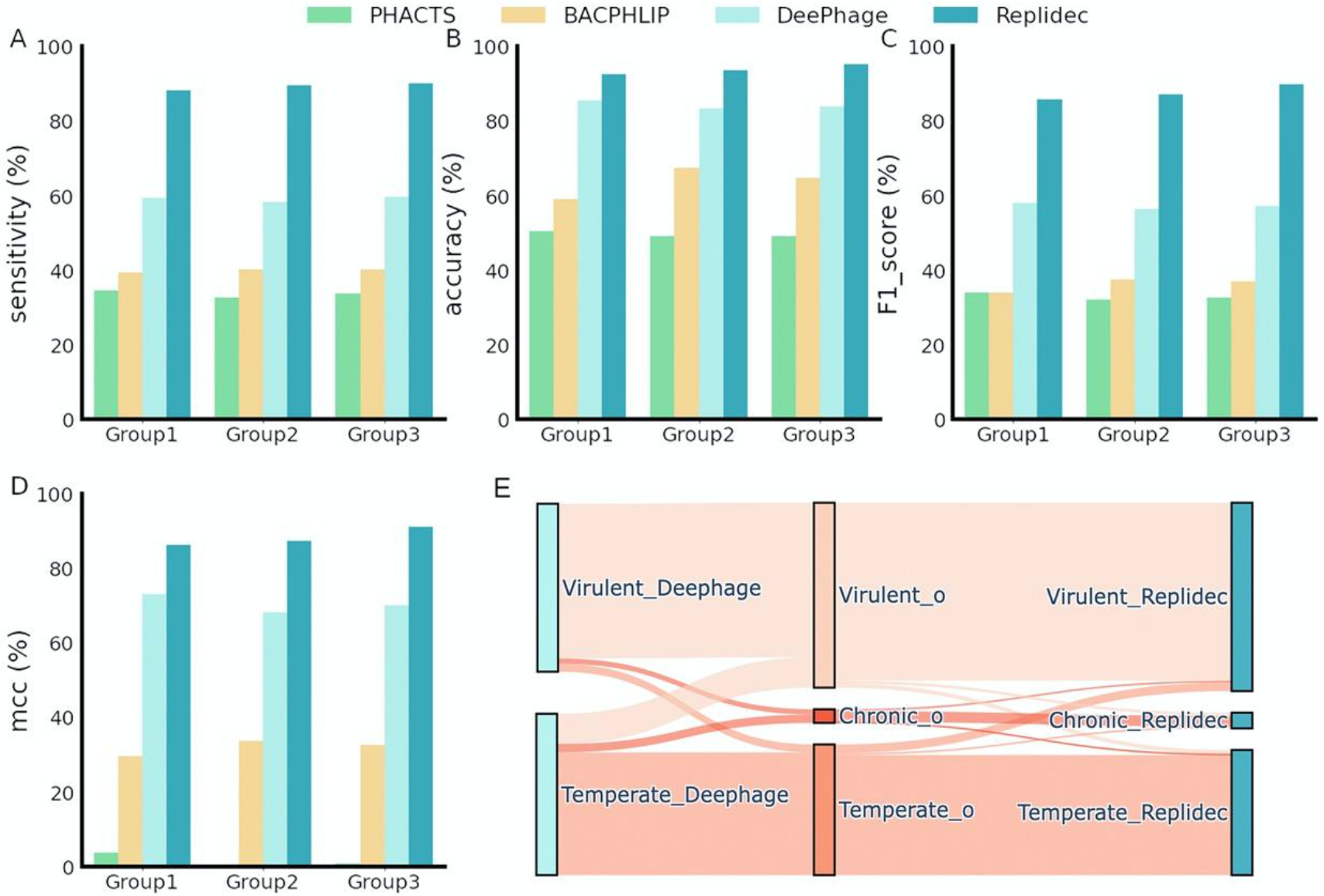
Performance on metagenomic assemblies. Comparison between four softwares on sensitivity(A), accuracy(B), f1_score(C), mcc(D). (E) detailed comparison between Deephage(left bar), Replidec(right bar) and original lifecycle of assemblies(middle bar with o as suffix).

We use the sankey plot to compare the prediction of DeePhage, and Replidec. Contrary to Deephage that inaccurately predicted the replication cycle of the 257 sequence from the 1.57k virulent sequences(Figure 3E) Replidec performed significantly better in predicting the replication cycle of viruses in the dataset.

We also tested how input sequence length influences the prediction(Figure S3). Bacphlip and Deephage perform well on very long query sequences, and their predictions were improved by increasing the length of genomes while Replidec predictions were not influenced by the query length.

## Discussion

Phages are believed to play a significant role in most ecosystem by regulating the bacterial abundance, diversity, and metabolisms [5]. Phage mainly replicated through three replication cycles: lytic, lysogenic, and chronic cycles [5]. Multiple tools have recently been developed for prediction of replication cycle in viral sequences. However, these tools have already provided valuable information about the replication cycle of viruses. They have limitations with their prediction. Here, we present Replidec, which uses alignment-based approach (relaying on marker genes such as Integrase, excisionase, PI-like genes for inoviruses) and naïve bayes classifier (for sequences with no marker genes) to predict replication cycle for viral sequences.

In addition, most in silico tools developed so far classify phages into two types of replication cycles (lytic and lysogenic). This is because these softwares use training datasets which do not contain genomes of phages that replicate through chronic lifecycle, such as PHANTOME database(www.phantome.org), which is used by PHACT and Deehage. PhagePred uses the dataset from Mavrich[20] and BACPHLIP uses the strategy developed by Mavrich[20]. PhageAI use PhagesDB[21] (https://phagesdb.org/clusters/) and ACLAME[22] (http://aclame.ulb.ac.be). Since inoviruses are found in every most microbial habitat and they are globally distributed[7], predictions of replication cycles by these tools are questionable. Therefore, developing new tools are essential for reliably predicting the replication cycles in virome data. In contrast, Replidec can predict all three replication cycles in viral sequences with a high accuracy.

Yet, Replidec also has limitations. For example, like other alignment-based softwares which rely on the databases for their predictions Replidec also suffers from biases that exist in databases. The NCBI Refseq dataset might lack virus sequences from extreme environments. Furthermore, if query data has no or very few “hits” with the custom database were used, Replidec hardly makes a confidential decision. In order to avoid this, we recommend that sequences with very few coding genes should not be used as inputs. We expect this to be improved with more genomes available in public databases. In addition, since the naïve bayes classifier used by Replidec is assuming that each protein feature is independent while in real circumstances, some genes may work together in biological processes[24][25]. For example, CI, CII, and Cro, in phage λ, control lysis-to-lysogeny decision[26]. However, Replidec is not using this information in its predictions since it is difficult to measure these types of interactions in mathematical models plus they are not fully understood.

## Conclusions

Replidec use a naïve bayes classifier to predict viruses replication cycle, especially for the metagenomics derived sequences. Replidec can achieve better performance in complete genomes and in simulated metagenomic data compared to most prediction softwares (except phageAI).

## Methods

### Datasets

#### Custom viral protein database (Viral_Protein_DB)

We made an in house database (Viral_Protein_DB) by acquiring viral genomes and bacterial genome from NCBI (download: 2021-09-21 and 2021-10-27). Viral genome acquired from Refseq Database and Bacteria genomes accessions were fetched from NCBI (https://ftp.ncbi.nlm.nih.gov/genomes/refseq/assembly_summary_refseq.txt) by setting the ‘RefSeq category’ equal to ‘reference genome’ and ‘representative genome’.

First, we predicted all viral CDSs from viral genomes and prophages within bacteria genomes, respectively. CDSs from viral genomes are predicted using Prodigal V2.6.3[27]. Prophages and prophages CDSs within bacterial genomes are predicted by VIBRANT v1.2.1 [29]. We only pick prophage CDSs from a temperate prohages (predicted by VIBRANT) and its amino acid length greater than 80 amino acid. In total, 1,323,491 CDSs are retained, 584,295 CDSs were predicted from 14,717 viral genomes and 739,196 prophage CDSs were retained from 21,134 lysogenic prophages within 14,922 bacteria and archaea genomes. We merged amino acid sequences of these CDSs to build Viral_Protein_DB. Viral_Protein_DB were annotated based on HMM searches against 4 databases: Pfam(v34), kegg(download date: 2022-02-01), vog (211), Phrogs (http://millardlab.org/2021/11/21/phage-annotation-with-phrogs/) under 1e-5 using HMMER. Replication cycle of viral genomes is predicted using BACPHLIP and prophages are labelled as ‘Temperate’. Next, we used MMseqs2 [30] (Version: 7aade9df7475ae7c699b2074b5e4daa52e0245f1) with parameter “--cov-mode 0 --min-seq-id 0.70 -c 0.70”) to cluster the all amino acid sequences of all CDSs based on their identity of >70%. In total, 803,082 protein cluster are generated. We then caluculate the joint probablity for each protein cluster for naïve bayes classifer. And we create a gene sharing network of all genomes and PCs (cluster size greater than 15) using Cytoscape.

#### Testing Dataset

PhageAI website(https://app.phage.ai/) provide a detailed viruses information list and we pick the genome with known origin replication cycle and genome coverage is equal to “complete genome”. 1028 genomes were download based on the accession number from NCBI. The data were split into three groups (Group1, Group2, Group3) based on their diversity in replication cycle, phage family, and host. Each group includes over 300 phages (Group1:129 temperate, 136 virulent and 64 chronic; Group2: 113 temperate and 188 virulent and 62 chronic; Group3: 117 temperate and 155 virulent and 64 chronic;). And we use PHACT, BAPHLIP, PhageAI and Replidec to predict the replication cycle of these complete viral genomes. Then these genomes of each group is use to simulate the 150bp paired-end metagenomics reads by ART v2.5.8 [28] (parameter: -ss HS25 -p -l 150 -f 10 -m 200 -s 10).

#### Mathematic model of Replidec

The naïve Bayes Classifier technique[29] is based on Bayesian theorem and is particularly suited for high dimensional inputs. In order to reduce the complexity of this high dimensionality, the naïve Bayes classifier assumes that the features are mutually independent given a class. That is to say, one features is not affected by other feature.

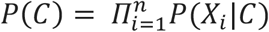

Where **X** = (*X*_1_, …, *X*_*i*_) means the feature vector and *C* is class. Here each protein cluster from cluster step is the feature vectors and *C* is the replication cycle type (Virulent or Temperate). Based on Bayesian theorem, it will be easy to calculate *P*(*X*).

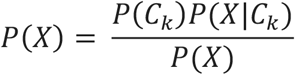

Because the denominator is a constant. So in practice, the formulation can be expressed as follow:

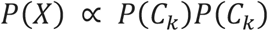

By using the chain rule, the finally formulation is,

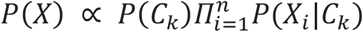

Where **X** = (*X*_1_, …, *X*_*i*_) means the feature vector and *C*_*k*_ means class. k = (0,1) means two type of replication cycle. *P*(*C*_k_) is the prior probablity and *P*(*C*_0_) is the number of temperature viruse genomes divided the total number of genome used in Viral_Protein_DB. *P*(*C*_1_) is the number of virulent viruses divided the total number of genome used in Viral_Protein_DB.

In order to avoid *P*(*C*_*k*_) equal to 0, a small number *α* was add to each feature vector (*α* = 1) given *C*_*k*_. And to reduce the computation complexity, base 10 log form was applied.

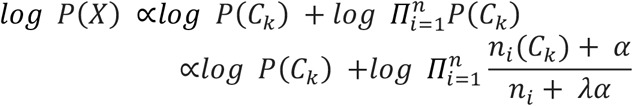

Where *n*_*i*_(*C*_*k*_) denotes the number of the ith feature vector in the k class and *λ* denotes the number of class (here *λ* = 2).

If the *log log P*((*X*_1_, …, *X*_*j*_)) > *log P*(*X*_1_, …, *X*_*j*_) (which *j* ≤ *n*), the lifestyle will be labed as “Temperate”, otherwise will be labeled as “Virulent”.

Based on the mathematical model, we generate a probability profile of each protein cluster in 2 lifestyle classes (temperate and virulent).

#### The Replidec pipeline

The Replidec pipeline is written in Python and some open source software were used: Prodigal[27], HMMER[30], MMseqs2[31], Blastp[32].

The Replidec software cover 3 steps(Figure 4):

**Figure 4:**
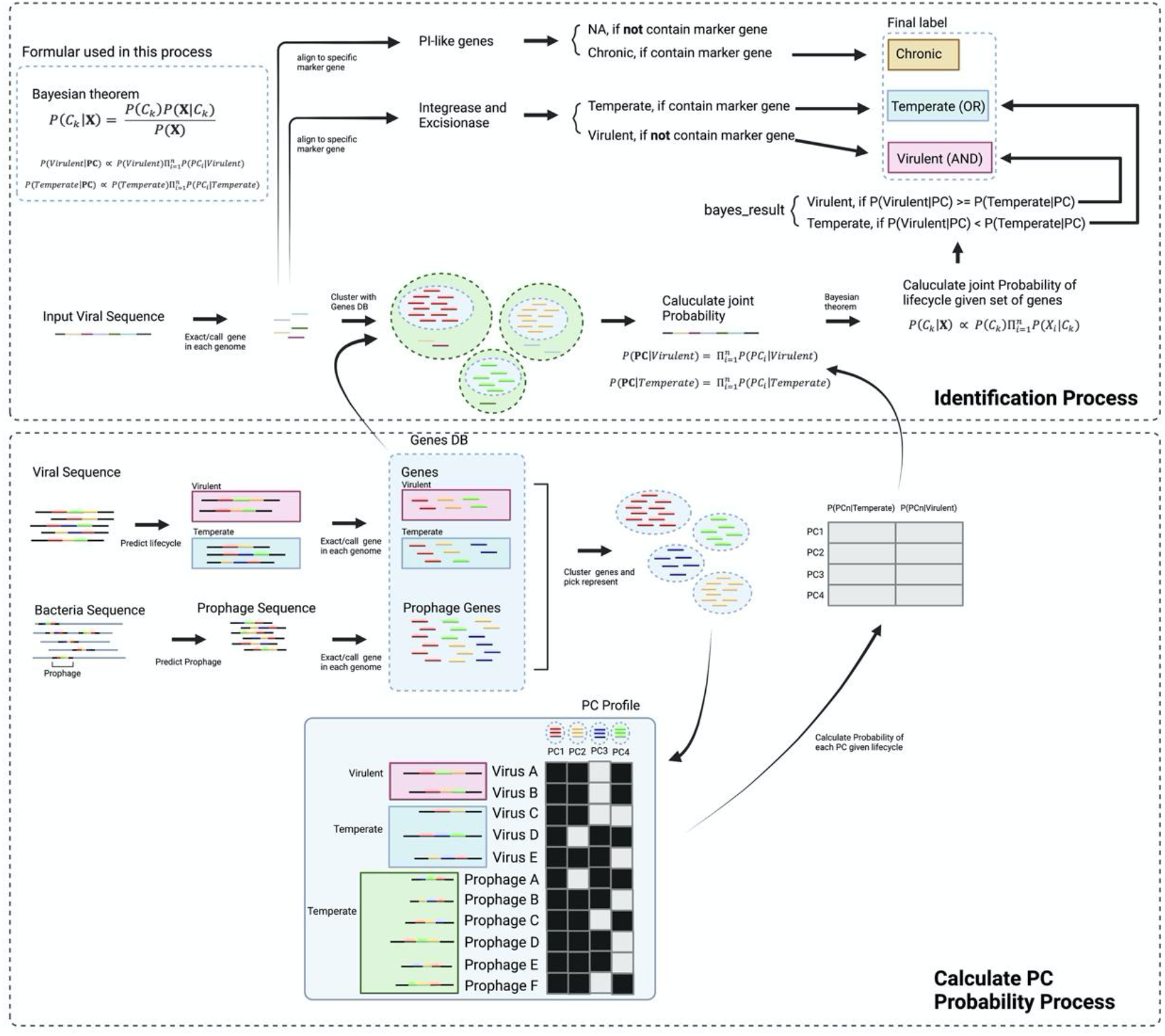
The Replidec pipeline: proposed methodology for virus replication cycle prediction.

1. Biological search for integrase and excisionase using hmmsearch[30]
  a. search integrase (contain 27 pfam family) and excisionase (contain 3 pfam family) existence in the query data.
  b. If they exist in the query data, the label of biological search will be labeled as “Temperate”.
2. Biological search for PI-like genes db using hmmsearch[30] and blastp[32], methods are from Roux’s article[7]
  a. If they exist in the query data, the label of biological search will be labeled as “Chronic”.
3. Computational calculation of lifestyle probablity using naïve bayes Classifier
  a. Align query data with 1,323,491 genes in Viral_Protein_DB using Mmseqs2 (easy-search, with paramter ‘-s 7 --max-seqs 1 --alignment-mode 3 --alignment-output-mode 0 --min-aln-len 40 --cov-mode 0 --greedy-best-hits 1’)
  b. calculate the conditional probablity of *P*(*X*), Where **X** = (*X*_1_, …, *X*_*i*_) means the maped protein cluster and *C*_*k*_ means class. k = (0,1) means two type of lifestyle.
  c. if the *log log P*(*X*) greater than *log log P*(*X*), the label of Computational calculation will be labeles as “Temperate”.

If one of 1 or 3 steps give a “Temperate” label, the final predicted result will be labeled as “Temperate”.

If the input data is a contig or genome then a preprocess step will be executed: predict the gene in the input file using prodigal (-g 11).

#### Simulated dataset workflow

To test the performance on the simulated metagenomics data. Simulated PE reads were followed a main analysis workflow. Simulated 150bp PE reads were using fastp to filter low quality reads with default parameter then using SPAdes v3.15.2 (--meta) [33] to assemble. Retain contigs length greater than 3000 bp. Total 2795 assemblies were generated and 882, 957, 956 contigs from Group1, Group2 and Group3, respectively. The assembled contig is compared with original genomes to find the most similar origin sequence using Minimap2[34]. PHACTS[11], BACPHLIP[12], DeePhage[15] were used to predict the contigs lifestyle. PhagePred is not include, because detail comparison was made in Deephage paper.

#### >Evaluation metrics

We use the criteria of Sensitivity (Sn), Accuracy (Acc), F1-score and Matthews correlation coefficient (Mcc) to evaluate different prediction methods, which are calculated as follows:

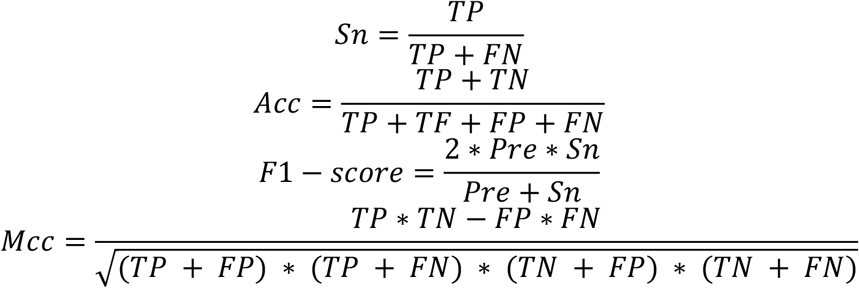

where TP, FN, TN and FP denote the numbers of true positive, false negative, true negative, and false positive, respectively. All the metrics are calculated using scikit package, average sensitivity and average F1-score of multiclass are calculated using “macro” parameter.

## Acknowledgments

This work was funded by the German Research Foundation (D.F.G. Emmy Noether program, Project No. 273124240, SFB 1371, Project No. 395357507), Marie Sklodowska-Curie Actions Innovation Training Networks grant agreement no. 955974(VIROINF), and the European Research Council Starting grant (ERC StG 803077) awarded to L.D.

## Author Contributions

X.P. and J.R. developed the software. X.P. constructed the database, performed the analyses, and drafted the manuscript. M.K.M. and L.D. conceived and supervised the project and drafted the manuscript. All authors reviewed and approved the manuscript.

## Reference

1. Microbiology by numbers. Nat Rev Microbiol. 2011;9:628–628.

2. Breitbart M, Thompson L, Suttle C, Sullivan M. Exploring the Vast Diversity of Marine Viruses. Oceanography. 2007;20:135–9.

3. Camarillo-Guerrero LF, Almeida A, Rangel-Pineros G, Finn RD, Lawley TD. Massive expansion of human gut bacteriophage diversity. Cell. 2021;184:1098-1109.e9.

4. Mirzaei MK, Maurice CF. Ménage à trois in the human gut: interactions between host, bacteria and phages. Nat Rev Microbiol. 2017;15:397–408.

5. Hobbs Z, Abedon ST. Diversity of phage infection types and associated terminology: the problem with ‘Lytic or lysogenic.’ FEMS Microbiol Lett. 2016;363:fnw047.

6. Clokie MR, Millard AD, Letarov AV, Heaphy S. Phages in nature. Bacteriophage. 2011;1:31–45.

7. Roux S, Krupovic M, Daly RA, Borges AL, Nayfach S, Schulz F, et al. Cryptic inoviruses revealed as pervasive in bacteria and archaea across Earth’s biomes. Nat Microbiol. 2019;4:1895–906.

8. Roux S, Adriaenssens EM, Dutilh BE, Koonin EV, Kropinski AM, Krupovic M, et al. Minimum Information about an Uncultivated Virus Genome (MIUViG). Nat Biotechnol. 2019;37:29–37.

9. Khan Mirzaei M, Khan MdAA, Ghosh P, Taranu ZE, Taguer M, Ru J, et al. Bacteriophages Isolated from Stunted Children Can Regulate Gut Bacterial Communities in an Age-Specific Manner. Cell Host Microbe. 2020;27:199-212.e5.

10. Clooney AG, Sutton TDS, Shkoporov AN, Holohan RK, Daly KM, O’Regan O, et al. Whole-Virome Analysis Sheds Light on Viral Dark Matter in Inflammatory Bowel Disease. Cell Host Microbe. 2019;26:764-778.e5.

11. McNair K, Bailey BA, Edwards RA. PHACTS, a computational approach to classifying the lifestyle of phages. Bioinformatics. 2012;28:614–8.

12. Hockenberry AJ, Wilke CO. BACPHLIP: Predicting bacteriophage lifestyle from conserved protein domains. :6.

13. Tynecki P, Guziński A, Kazimierczak J, Jadczuk M, Dastych J, Onisko A. PhageAI - Bacteriophage Life Cycle Recognition with Machine Learning and Natural Language Processing. preprint. Bioinformatics; 2020.

14. Song K. Classifying the Lifestyle of Metagenomically-Derived Phages Sequences Using Alignment-Free Methods. Front Microbiol. 2020;11:567769.

15. Wu S, Fang Z, Tan J, Li M, Wang C, Guo Q, et al. DeePhage: distinguishing virulent and temperate phage-derived sequences in metavirome data with a deep learning approach. GigaScience. 2021;10:giab056.

16. Shao Q, Trinh JT, Zeng L. High-resolution studies of lysis-lysogeny decision-making in bacteriophage lambda. J Biol Chem. 2019;294:3343–9.

17. Zhang L, Xu D, Huang Y, Zhu X, Rui M, Wan T, et al. Structural and functional characterization of deep-sea thermophilic bacteriophage GVE2 HNH endonuclease. Sci Rep. 2017;7:42542.

18. Hanley-Bowdoin L, Bejarano ER, Robertson D, Mansoor S. Geminiviruses: masters at redirecting and reprogramming plant processes. Nat Rev Microbiol. 2013;11:777–88.

19. Krupovic M, Ravantti JJ, Bamford DH. Geminiviruses: a tale of a plasmid becoming a virus. BMC Evol Biol. 2009;9:112.

20. Mavrich TN, Hatfull GF. Bacteriophage evolution differs by host, lifestyle and genome. Nat Microbiol. 2017;2:17112.

21. Russell DA, Hatfull GF. PhagesDB: the actinobacteriophage database. Bioinforma Oxf Engl. 2017;33:784–6.

22. Leplae R, Lima-Mendez G, Toussaint A. ACLAME: a CLAssification of Mobile genetic Elements, update 2010. Nucleic Acids Res. 2010;38 Database issue:D57–61.

23. Hatfull GF, Hendrix RW. Bacteriophages and their genomes. Curr Opin Virol. 2011;1:298–303.

24. Erez Z, Steinberger-Levy I, Shamir M, Doron S, Stokar-Avihail A, Peleg Y, et al. Communication between viruses guides lysis–lysogeny decisions. Nature. 2017;541:488–93.

25. Ofir G, Sorek R. Contemporary Phage Biology: From Classic Models to New Insights. Cell. 2018;172:1260–70.

26. Oppenheim AB, Kobiler O, Stavans J, Court DL, Adhya S. Switches in Bacteriophage Lambda Development. Annu Rev Genet. 2005;39:409–29.

27. Hyatt D, Chen G-L, LoCascio PF, Land ML, Larimer FW, Hauser LJ. Prodigal: prokaryotic gene recognition and translation initiation site identification. BMC Bioinformatics. 2010;11:119.

28. Huang W, Li L, Myers JR, Marth GT. ART: a next-generation sequencing read simulator. Bioinformatics. 2012;28:593–4.

29. Rish I. An empirical study of the naive Bayes classifier. :6.

30. Eddy SR. Accelerated Profile HMM Searches. PLoS Comput Biol. 2011;7:e1002195.

31. Steinegger M, Söding J. MMseqs2 enables sensitive protein sequence searching for the analysis of massive data sets. Nat Biotechnol. 2017;35:1026–8.

32. Johnson M, Zaretskaya I, Raytselis Y, Merezhuk Y, McGinnis S, Madden TL. NCBI BLAST: a better web interface. Nucleic Acids Res. 2008;36 suppl_2:W5–9.

33. Nurk S, Meleshko D, Korobeynikov A, Pevzner PA. metaSPAdes: a new versatile metagenomic assembler. Genome Res. 2017;27:824–34.

34. Li H. Minimap2: pairwise alignment for nucleotide sequences. Bioinformatics. 2018;34:3094–100.

