## Supplemental File for "Replidec - Use naive Bayes classifier to identify virus lifecycle from metagenomics data"

Supplement materia

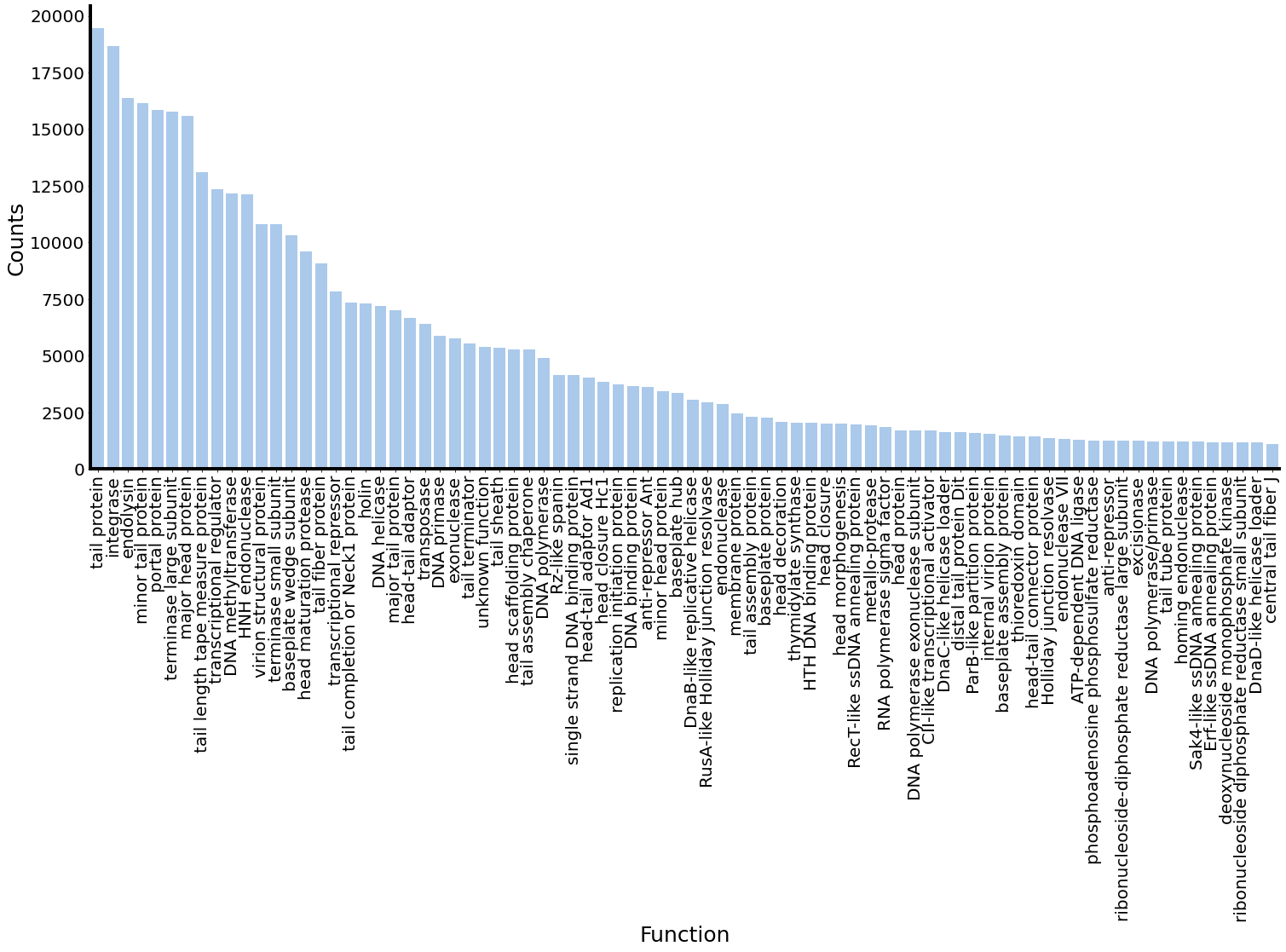

Figure S1: function distribution of Viral_Protein_DB

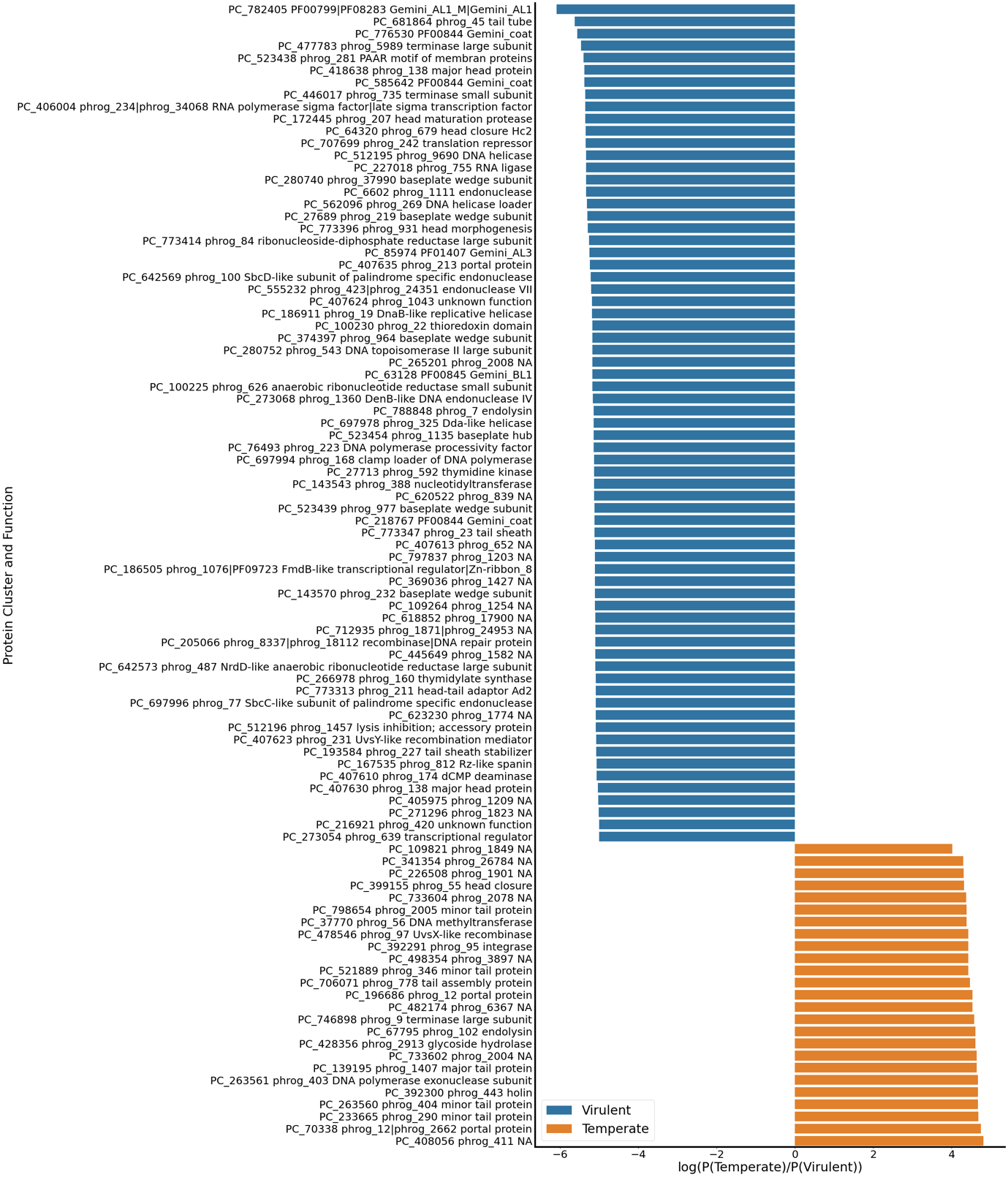

Figure S2: function distribution of Viral_Protein_DB protein cluster. (P(temperate) – P(Virulent)) is great than 4 and small than -5

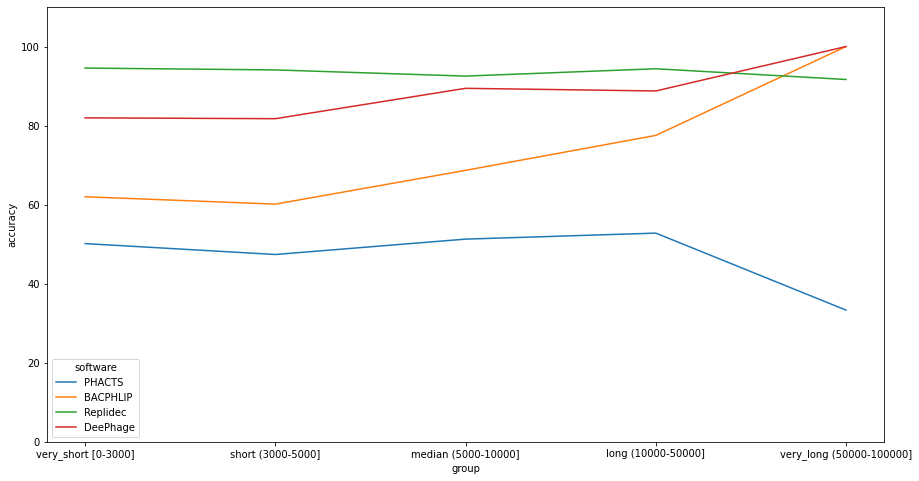

Figure S3: Length of simulated assemblies on predicting perfermance. Accuracy calculated using scikitlearn.
